# Interactome of vertebrate GAF/ThPOK reveals its diverse functions in gene regulation and DNA repair

**DOI:** 10.1101/559641

**Authors:** Avinash Srivastava, Rakesh K Mishra

## Abstract

Evolutionary conservation and lineage-specific diversification of existing proteins forms the basis for evolving complexity of protein-protein interaction networks and presumably, thereby that of organism. GAGA associated factor (GAF) belongs to BTB/POZ and zinc finger family of transcription factors and it is conserved from flies to humans. Emerging evidence shows indispensable roles of vertebrate GAF (vGAF a.k.a. ThPOK) in functionally divergent developmental processes like hematopoiesis, adipogenesis and lactation. vGAF is a sequence-specific DNA binding transcription factor with multiple context-dependent roles in gene activation/repression, enhancer-blocking and more recently, it has shown to be the part of ribonucleoprotein complexes as well. In order to understand the molecular basis of these diverse functions, we analyzed the protein-protein interactome of vGAF. This analysis shows vGAF’s association with chromatin remodelers, RNA metabolic machinery, transcriptional activators/repressors, and components of DNA repair machinery, thereby provides a plausible explanation for the diverse molecular functions of vGAF. Our findings discern novel role of vGAF in several molecular processes like DNA repair and RNA metabolism. We further tested the biological significance of our protein-protein interaction data and show a novel function of vGAF in DNA repair and cell-survival after UV induced DNA damage. Consistent with these results, analysis of high-throughput RNA-seq data shows the downregulation of vGAF in samples of skin cutaneous melanoma for which the primary cause is UV induced DNA damage. These findings suggest vGAF as an early diagnostic biomarker for skin cutaneous melanoma. Taken together, our study reveals a molecular basis for the diverse functions of vGAF. We uncover its role in DNA repair and provide an explanation for key roles of such evolutionary conserved factors in processes like development and disease.

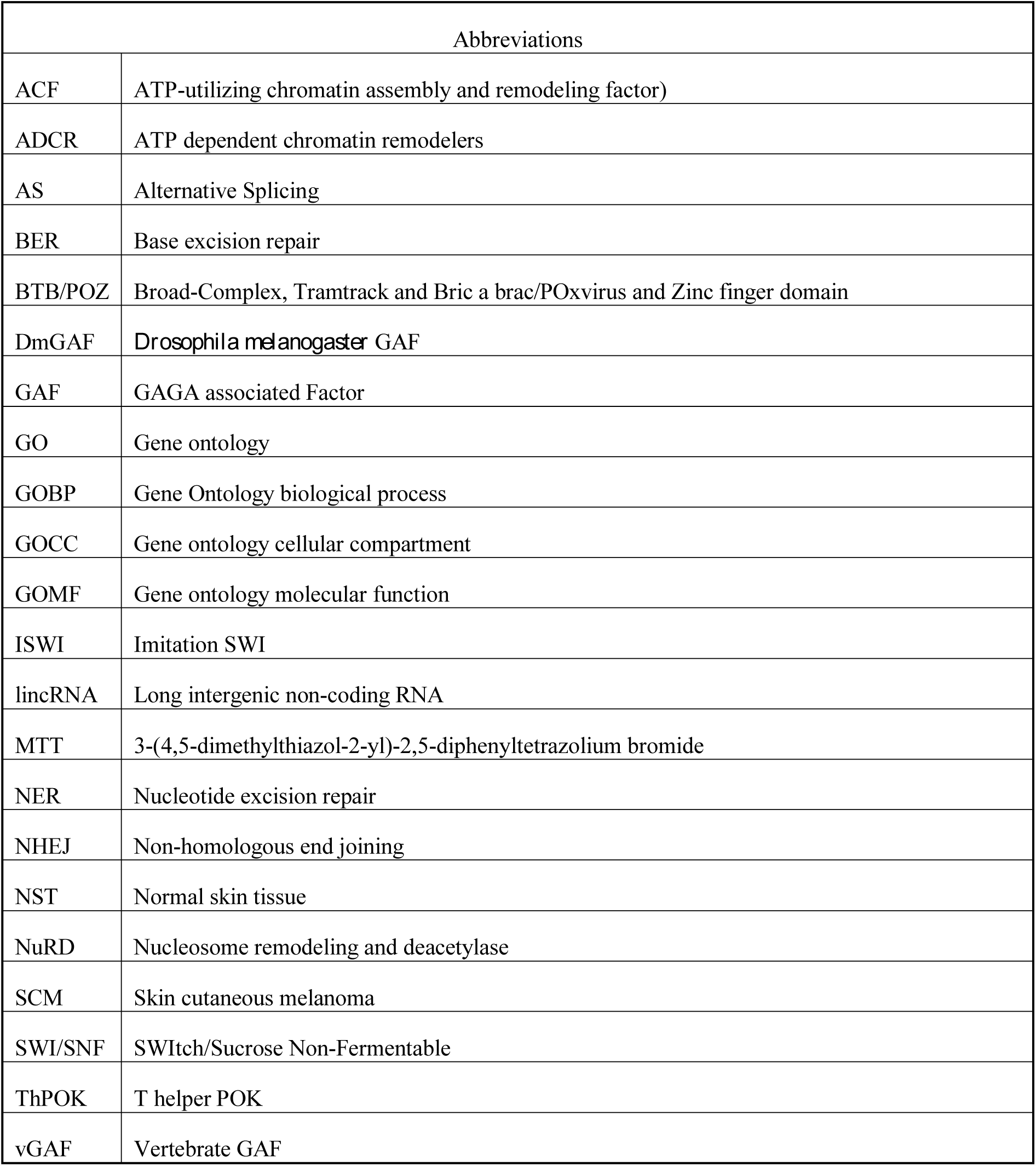

## 1. Introduction

GAGA associated factor (GAF) of *Drosophila melanogaster*, DmGAF, is a developmentally important transcription factor that has been implicated in diverse nuclear processes like gene activation, polycomb-mediated silencing, enhancer-blocking, position effect variegation and chromosomal segregation [1-6]. Vertebrate GAF (vGAF), the evolutionarily conserved counterpart of DmGAF in vertebrates, is encoded by *zbtb7b* gene [7]. ZBTB7B/vGAF/ThPOK has been shown to be essential for the proper development of the hematopoietic and adipose tissues [8-10]. A recent study also points out the role of vGAF in the onset of lactation and in the production of milk lipids [11]. The molecular functions of vGAF that translate into these biological processes are diverse in nature. vGAF (ZBTB7B/ThPOK) was first identified as a tissue-specific activator of collagen genes in mouse dermis [12,13]. Several other studies also point the role of vGAF in transcriptional activation of genes involved in cytokine signaling like *Socs1, Cish* and *TNF-*alpha [14,15]. Paradoxically, multiple other studies report that vGAF has a rather repressive effect on the transcription of another set of genes like *UDP glucose dehydrogenase, eomesodermin*, and *cd8* [16-18]. These are not, however, the only nuclear processes where vGAF has been shown to be involved. vGAF has been reported to bind at the enhancer-blocker elements in the murine *Hox* clusters and is essential for tissue-specific nuclear-lamina proximal positioning of *IgH* and *cyp3a* loci, suggesting an architectural role of vGAF in chromatin organization [19,20]. Moreover, a recent study has shown a developmentally important association of vGAF with ribonucleoprotein complexes containing a lincRNA (long inter-genic non-coding RNA) [10]. The remarkable functional versatility of vGAF, at least in part, may stem from the structural domains of the protein [21]. The zinc-finger domains present at C-terminal of vGAF allow sequence-specific tethering of vGAF molecules at numerous loci in the genome, while, the N-terminal BTB/POZ domain being a protein-protein interaction domain endows GAF with an ability to recruit protein-complexes of diverse functionalities at these loci. Several pieces of evidence suggest the importance of protein-protein interactions in the multifunctional roles of GAF proteins [16,20,22-24]. We hypothesize that like DmGAF, the functional versatility of vGAF depends on its interacting partners, which themselves are functionally diverse.

In the present study, we sought to decipher a comprehensive protein-interaction network of vGAF. We used mass spectrometry and identified 585 proteins that co-immunoprecipitate with vGAF. These proteins belong to diverse functional classes, viz., chromatin remodelers, transcriptional activators/repressors, RNA processing factors and components of DNA repair machinery. We show that vGAF co-localizes to the site of DNA damage and further establish an important role of vGAF in DNA repair and cell-survival after UV induced DNA damage. The diversity of the vGAF-associated protein complexes observed in this study explains the molecular basis for the diverse functions of this protein, and also implicate its function in multiple other nuclear processes.

## 2. Material and Methods

### 2.1 Plasmids and antibodies

ThPOK cDNA (IMAGE ID-6309645) was obtained from Open-Biosystems (Dharmacon). The N-terminal 3X-FLAG-tagged ThPOK expression plasmid was generated by cloning the ThPOK coding DNA sequence (CDS) in the backbone of pEGFPC-3 (Clontech) plasmid after replacing the CDS of EGFP with 3X-FLAG. CDS of MBD3, Cbx5, HDAC1, p65, and RBM14 were amplified from C2C12 myoblast cell cDNA using Q5® High-Fidelity 2X Master Mix. Expression plasmids of EGFP-tagged MBD3, Cbx5, HDAC1, p65, and RBM14 were generated by cloning the respective CDS of proteins into the pEGFPC-1 plasmid (See Sup. Table 3 for primer and restriction site details).

For western blotting, anti-FLAG M2 mouse monoclonal antibody was procured from Sigma and anti-GFP (ab290) rabbit polyclonal was from Abcam. p-Histone H2A.X (Ser-139)/γ-H2A.X (SC-10196) was obtained from Santacruz. Secondary antibodies anti-Mouse HRP (ab6820) and anti-Rabbit HRP (ab6802) were obtained from Abcam. For immuno-staining, p-Histone γ-H2A.X (Ser-139) (clone JBW301, cat. 05-363) was from Millipore while anti-FLAG rabbit polyclonal (F7425) antibody was procured from Sigma. Fluorochrome-conjugated secondary antibodies anti-Rabbit IgG (H+L) Alexa-flour 647 (cat. 711-605-152) and anti-Mouse IgG (H+L) Cy3 (cat. 115-165-062) were obtained from Jacksons laboratories.

### 2.2 RNA isolation and cDNA preparation

Total RNA was prepared from C2C12 myoblast cells using TRIzol (Ambion, Invitrogen) following the manufacturer’s instructions. One microgram of total RNA was used for preparing cDNA using PrimeScript™ 1st strand cDNA Synthesis Kit (Clontech) following the manufacturer’s protocol.

### 2.3 Cell culture and transfection

C2C12 myoblast cells were maintained in Dulbecco‘s modified Eagle‘s medium (DMEM) supplemented with 20% fetal bovine serum (FBS) and 1X GlutaMax (Invitrogen). HEK293 cells were maintained in DMEM supplemented with 10% FBS. C2C12/HEK293 cells were grown in 100mm tissues culture dishes and were transfected with 12ug plasmid DNA using Lipofectamine-LTX (C2C12 cells) or Lipofectamine-3000 (HEK293) following the manufacture’s instruction. Primary fibroblast cell cultures were established using depilated skin and lung tissue of vGAF knockout mice (The Jacksons laboratory: Stock No:027663) and wild type sex-matched littermates using a previously published protocol [8,25]. All cell cultures were maintained at 37° C with 5% CO_2_ in a humidified chamber.

### 2.4 Immuno-precipitation, protein-extraction and western blotting

C2C12 myoblast/HEK293 cells were harvested 36 hours post transfection and washed twice with ice-cold PBS. Cells were resuspended in cell-lysis buffer (50mM Tris pH 7.4, 150mM NaCl, 0.5mM MgCl_2_, 1% Triton-X 100, 10% glycerol, 1X protease inhibitor cocktail (Roche), 1mM PMSF, 270U/ml DNAase-I (sigma)) and incubated on ice for 45 minutes with intermittent mixing. The cell debris were removed by spinning the cell lysate at 14000rpm/4 °C for 10 min. ANTI-FLAG M2 (Sigma) mouse monoclonal or anti-GFP rabbit polyclonal (ab290) antibody was incubated with Protein-G/A magnetic beads (Dynabeads, Invitrogen) in PBST for 2hrs at 4 °C. Subsequently, Antibody-Protein-G/A magnetic bead complex was incubated with total cell lysate at 4 °C for overnight. The immune-complexes captured on beads were washed thrice with cell-lysis buffer (without DNAase-I) and eluted in the laemmli buffer.

Protein extracts were prepared from skin explants or primary cells. Skin explants were pulverized in liquid nitrogen. Pulverized skin or cells were resuspended in protein-extraction buffer (50mM Tris pH 7.4, 150mM NaCl, 1% Triton-X 100, 1% SDS, 10% glycerol, 10mM DTT, 1X protease inhibitor cocktail (Roche), 1mM PMSF) and incubated on ice for 45 minutes with intermittent mixing. The cell and tissue debris were removed by spinning the extracts at 14000rpm/4° C for 10 min. Protein samples were resolved on SDS-PAGE and transferred onto PVDF membrane followed by antibody-mediated detection using enhanced chemiluminescence method.

### 2.5 UV treatment

UV treatments were carried out using CL-1000 UV cross-linker (UVP). Exponentially growing cultures of primary cells were exposed to UVC (254nm) at a dose of 5J/m^2^ (lung cells) or 30J/m^2^ (skin cells). C2C12 cells were grown on glass coverslips and 36 hours after the transfection were UVC treated at a dose of 5J/m^2^. Depilated skin from vGAF knockout and wild type mice were cut into 1mm^2^ size and were kept with their dorsal side up in 6 well tissue culture plate. Skin explants were exposed to UVC at a dose of 60J/m^2^. Prior to protein extract preparation, tissues and cells were allowed to recover for 3hrs in DMEM medium supplemented with 10% FBS.

### 2.6. MTT assay

Skin primary cells were UV treated in 24 well tissue culture plates after 1-day post seeding at a density of 5000 cells/well. Cells were allowed to grow for another 96 hours and were washed with PBS before adding 100ul of MTT solution (0.5mg/ml MTT in PBS) to each well. Cells were incubated with MTT solution for another 4 hours before adding 150ul of DMSO to each well for the solubilization of farmazan crystals. Finally, the absorbance was measured at 570nm using using Multiskan microplate photometer (Thermo Scientific, USA).

### 2.7 Immuno-staining and microscopy

Cells were grown over glass coverslips and fixed with 4% paraformaldehyde. Next to the fixation, cclls were permeablized in PBST (0.5% Triton-X100) and were kept in blocking solution (1X PBS, 0.1% Tween-20, 1%BSA) for 1 hour. Cells were incubated with anti-γ-H2AX (1:1000) and anti-FLAG (1:1000) antibodies for 1 hour at room temperature. Fluorochrome-conjugated secondary antibodies used for subsequent detection of primary antibody binding were Anti-Rabbit IgG (H+L) Alexa-flour 647 (1:1000) and anti-Mouse IgG (H+L) Cy3 (1:500). Finally, coverslips were mounted on glass slides using Vectashield™ mounting media (with DAPI).

The optical sections of immune-stained cell nuclei were captured using Ziess LSM 880 confocal microscope with 63X objective at 3X zoom. Single plane optical sections of 5um thickness were used for co-localization analysis using ‘Colocalization Finder’ plugin of ImageJ.

### 2.8 Mass spectrometry and protein identification

Immuno-precipitated samples were fractioned by 12% SDS-PAGE. The gels were stained with Imperial™ protein stain (Thermo-Scientific), destained and washed with Milli-Q water several times. Each gel lane was sliced into six pieces and these gel-pieces were trypsin digested using a previously published protocol [26]. The resulting tryptic peptides were resolved using reverse phase chromatography. 18ul of the sample was loaded on reverse phase C18-Biobasic PicoFrit™ chromatography columns using Easy nLC-II (Thermo-Scientific) with a flow rate of 0.4ul/min; peptides were eluted on 60 min gradient (3% to 70% acetonitrile). Chromatographically separated peptides were analyzed using Q-exactive mass spectrometer. The top 10 peptide precursors were selected for MS/MS analysis. The mass spectra obtained were searched against mouse proteome from UniProt using SEQUEST HT algorithm incorporated in Thermo Proteome Discoverer (version 1.4.0.288). Enzyme specificity was set to full trypsin digestion with maximum 2 missed cleavages. Precursor mass tolerance was set to 10ppm and fragment mass tolerance was 0.6Da. Peptide identifications were accepted only if they pass the following criteria: Maximum Delta Cn value - 0.1, Minimum Xcorr value – 1.92, peptide confidence – High. Proteins identifications were accepted only if they cross the Percolator score −10. We identified a non-redundant and cumulative list of 853 proteins from three FLAG-vGAF pull-downs while two negative control pull downs yielded a total of 935 non-redundant proteins. We removed the proteins that were common between vGAF pull down and negative control pull down to identify 585 specific interactors of vGAF.

### 2.9 Gene ontology analysis of protein interactome

List of protein interactors was analysed using Cytoscape 3.5.1 and its plugin ClueGO [27,28]. In order to identify the functionally enriched clusters of Gene ontology (GO) terms, we performed enrichment analysis of GO terms (Biological process, Molecular function, Cellular compartment) using two-sided hypergeometric test with Bonferroni step down p-value correction. We used GO term fusions, and terms were considered significant only at p=<0.01. The resultant significant terms were visualized in an organic network layout where nodes were connected above 0.4 kappa threshold.

### 2.10 Bioinformatic analysis of gene expression data

We used Expression-DIY module of GEPIA (Gene Expression Profiling Interactive Analysis) to analyze RNA-seq data for skin cutaneous melanoma (SCM) and normal skin tissue (NST) from TCGA (The Cancer Genome Atlas) and GTEx (Genotype-Tissue Expression) databases. Box-plot function was used for plotting the differential expression of genes in SCM and NST samples using default settings. We used the stage-plot function for plotting the expression of vGAF across major stages of skin cutaneous melanoma using default settings

## 3. Result

### 3.1 Identification of vGAF associated protein interactome

To define the protein interactome of vGAF, we used immuno-affinity based purification of protein complexes and subsequently identified the proteins using liquid chromatography coupled with mass-spectrometry (LC-MS/MS). We tagged vGAF with an N-terminal 3X FLAG-tag and expressed the recombinant protein in C2C12 myoblast cells. Immuno-staining of FLAG-vGAF transfected cells with anti-FLAG antibody shows a very specific nuclear localization of FLAG-vGAF (Sup. Fig. 1). The whole cell lysate was used to immuno-precipitate the vGAF containing protein complexes using a monoclonal anti-FLAG antibody or a non-specific IgG (Fig. 1). Whole cell lysate prepared from empty-vector transfected C2C12 cells was also used for immuno-precipitation using the anti-FLAG antibody to serve as the negative control in the mass-spectrometry experiment. Immuno-precipitation experiments were done in triplicates (FLAG-vGAF transfected cells) or in duplicates (empty-vector transfected cells). Finally, the immuno-precipitated protein samples were analyzed using LC-MS/MS. Altogether, we could identify 585 proteins that specifically interact with vGAF across our three replicates (Sup. Table 1). Although, we have used the total cell extracts for all our experiments, cellular component ontology for these 585 proteins showed a statistical overrepresentation of nuclear proteins, suggesting the specificity of FLAG-vGAF immuno-precipitation (Sup. Fig. 2). We further validated our mass spectrometry data by reverse co-IPs followed by western blots. To this end, we selected five candidate proteins from the list of vGAF protein interactome viz. HDAC1, MBD3, (both are the parts of NuRD chromatin remodeling complex) [29], RBM14 (a protein that is involved in both DNA double stand break repair and transcription coupled alternative splicing) [30,31], CBX5 (Protein associated with heterochromatin) [32] and p65 (a context dependent transcription factor) [33]. These proteins were N-terminally GFP-tagged in pEGFPC-1 plasmid and were co-transfected with 3X-FLAG-tagged vGAF plasmid in HEK293 cells (Sup. Fig. 3a & 3b). Immuno-precipitation of GFP-tagged candidate proteins from co-transfected HEK293 cells shows enrichment of vGAF compared to IgG controls suggesting an *in-vivo* interaction between vGAF and candidate proteins (Fig. 2a & 2b). These results validate the efficacy of our experiment and identification of novel interacting partners of vGAF.

**Fig. 1.**
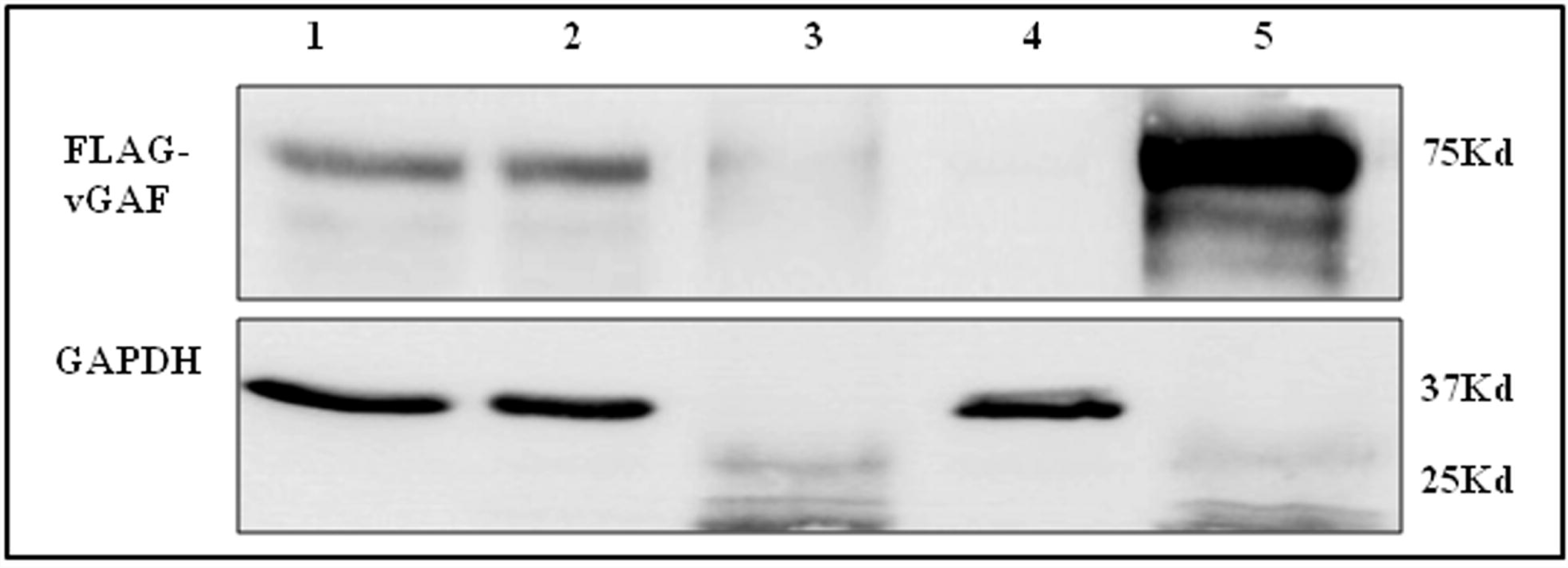
Immuno-precipitation of FLAF-vGAF. Western blot analysis of immuno-precipitated samples. FLAG antibody was used to assess the efficiency of the pull downs. Blots were probed with GAPDH antibody to check for non-specificity. Lane-1 – total cell extract (Input), Lane-2 – unbound fraction (output) after IP using IgG, Lane-3 – Immuno-precipitate using IgG, Lane-4 – unbound fraction (output) after IP using FLAG antibody, Lane-5 – Immuno-precipitate using FLAG antibody.

**Table 1.**
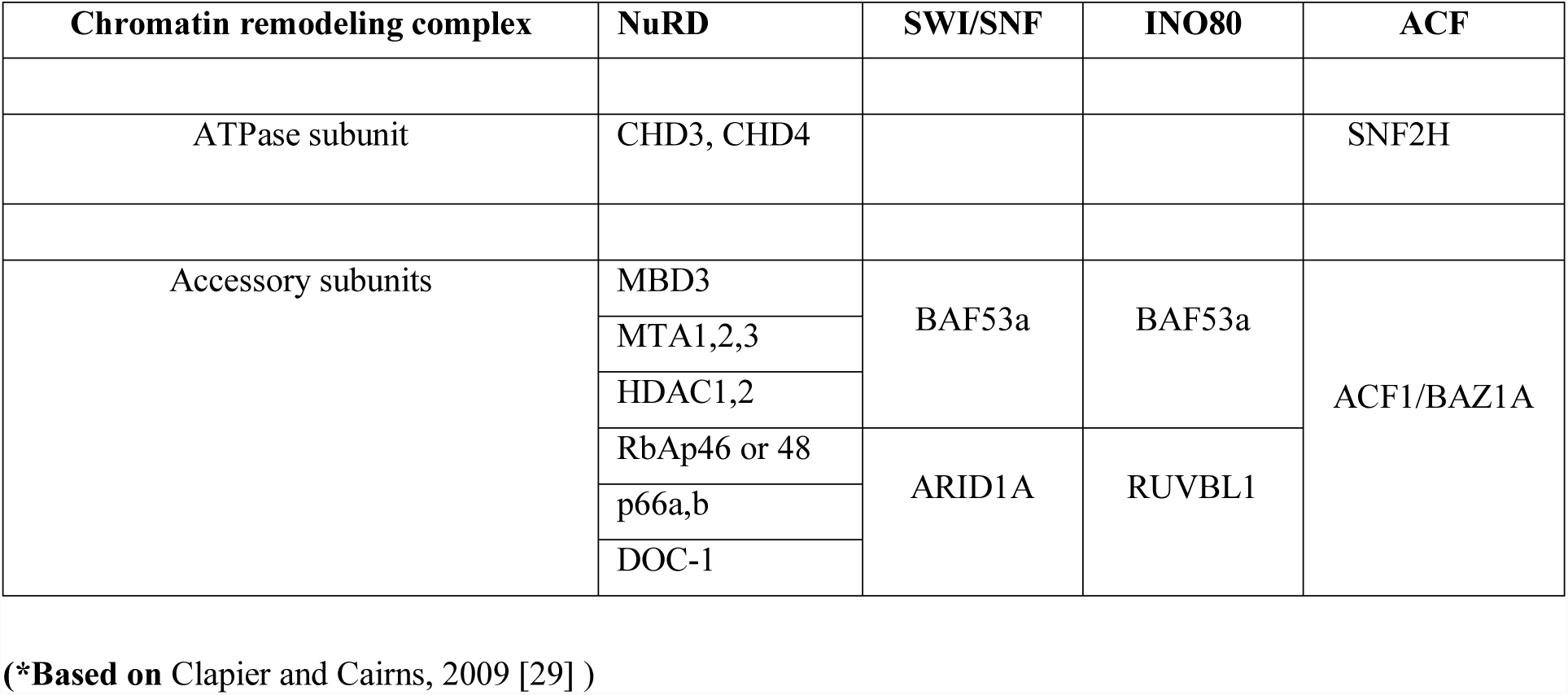
List of chromatin remodelers components* identified in vGAF protein interactome.

| Chromatin remodeling complex | NuRD | SWI/SNF | INO80 | ACF |
| --- | --- | --- | --- | --- |
| ATPase subunit | CHD3, CHD4 |  |  | SNF2H |
| Accessory subunits | MBD3 | BAF53a | BAF53a | ACF1/BAZ1A |
|  | MTA1,2,3 |  |  |  |
|  | HDAC1,2 |  |  |  |
|  | RbAp46 or 48 | ARID1A | RUVBL1 |  |
|  | p66a,b |  |  |  |
|  | DOC-1 |  |  |  |

(\*Based on Clapier and Cairns, 2009 [29] )

**Fig. 2.**
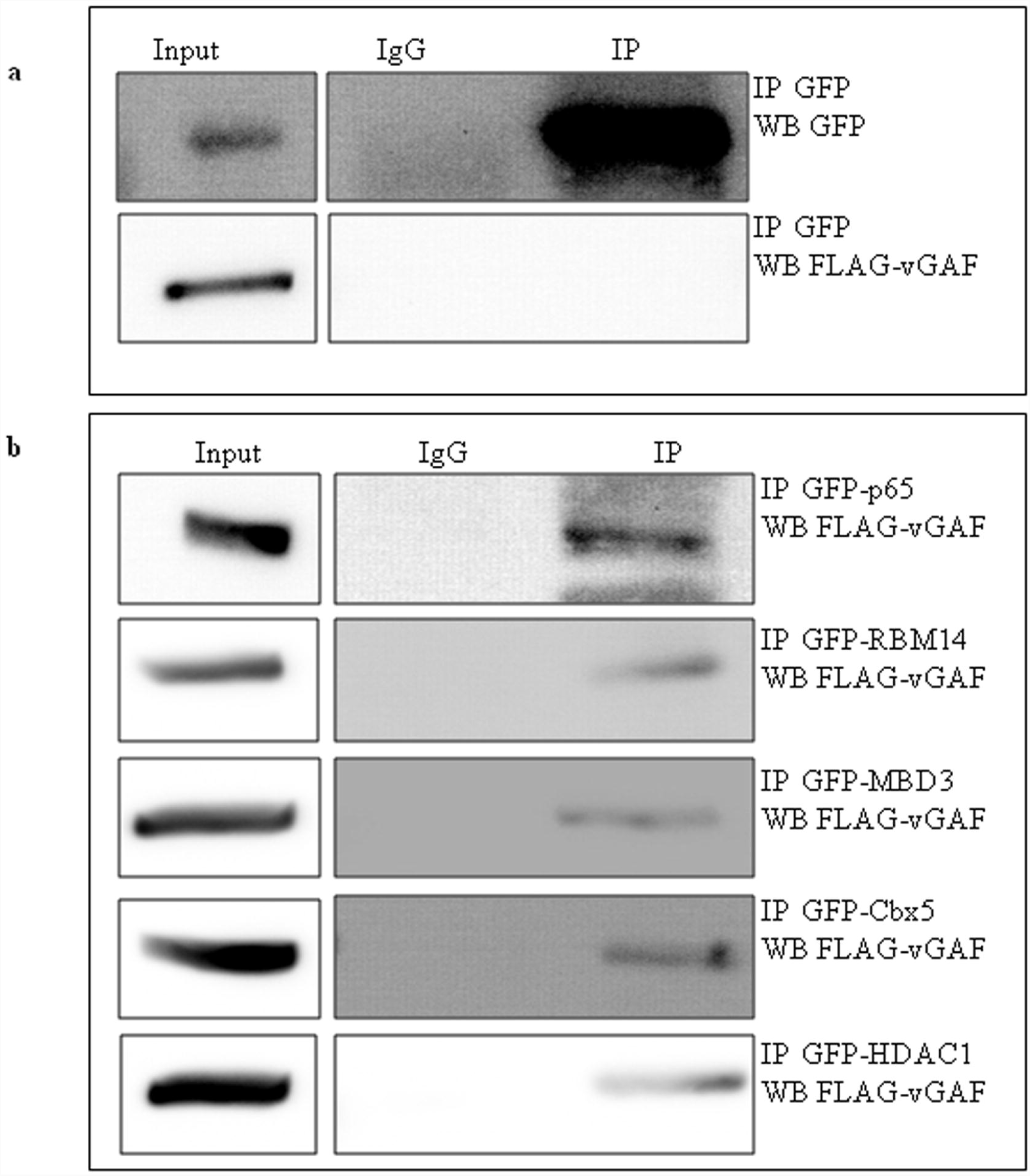
Validation of IP proteins with reverse co-IP. [a] FLAG-vGAF protein does not interact with EGFP. Immuno-precipitation (IP) was done using GFP antibody from HEK293 cells co-transfected with FLAG-vGAF and EGFP expression plasmids. Western blots were done with either anti-GFP antibody or anti-FLAG antibody. [b] FLAG-vGAF immuno-precipitates with N-terminal GFP tagged p65, RBM14, MBD3, Cbx5 and HDAC1. HEK293 cells were co-transfected with FLAG-vGAF and one of the N-terminal GFP tagged proteins (p65, RBM14, MBD3, Cbx5, HDAC1) constructs. IP was done with anti-GFP antibody and subsequently anti-FLAG antibody was used to detect the presence of FLAG-vGAF in immuno-precipitate using western blot (WB).

### 3.2 Gene ontology analysis of vGAF protein interactome

Gene ontology analysis of the vGAF protein interactome enabled us to identify the functional diversity among vGAF interacting partners. Fig. 3 shows a map of significantly enriched Gene ontology (GO) terms associated with vGAF interactome. The same results with additional details on genes associated with each GO term and statistical significance are presented in Sup. Table 2. The most significant Biological process associated GO terms (GOBP) in our analysis are ‘DNA metabolic process,’ ‘DNA Repair’ and ‘DNA Replication.’ Consistent with this we also find cellular component associated GO terms (GOCC) ‘Replication fork’ and ‘DNA replication factor C complex,’ suggesting a novel interaction of vGAF with components of DNA replication and repair. Another set of GOBP terms like ‘regulation of cellular macromolecule biosynthetic process,’ ‘nucleic acid templated transcription’, ‘transcription from RNA polymerase II promoter’ and ‘positive/negative regulation of gene expression’ etc. delineate the association of vGAF with RNA-polymerase-II associated macromolecular transcriptional complexes. Several other Molecular functions associated GO terms (GOMF) like ‘core promoter sequence-specific DNA binding’ and ‘sequence-specific double-stranded DNA binding’ also indicate the interaction of vGAF with other sequence-specific DNA binding proteins. GO analysis also shows the presence of GOMF terms ‘DNA-dependent ATPase activity’, ‘ATPase activity, coupled’ and GOCC term ‘SWI/SNF superfamily-type complex,’ suggesting the association of vGAF with ATP dependent chromatin remodelers. Interestingly, our analysis suggests the interaction of vGAF with components of RNA metabolic machinery through the presence of GOBP terms ‘RNA metabolic process’, ‘regulation of RNA metabolic process,’ ‘ribonucleoprotein complex assembly’ and GOCC term ‘spliceosomal complex’. In addition, our results show the presence of several unexpected GO terms viz. ‘Positive regulation of viral process,’ ‘Regulation of viral transcription,’ ‘ESCRT III complex,’ ‘cAMP response element binding’ and ‘Positive regulation of translation’ etc. Collectively, GO analysis of vGAF protein interactome clearly shows that vGAF interacts with protein complexes of diverse functionalities and nature.

**Fig. 3.**
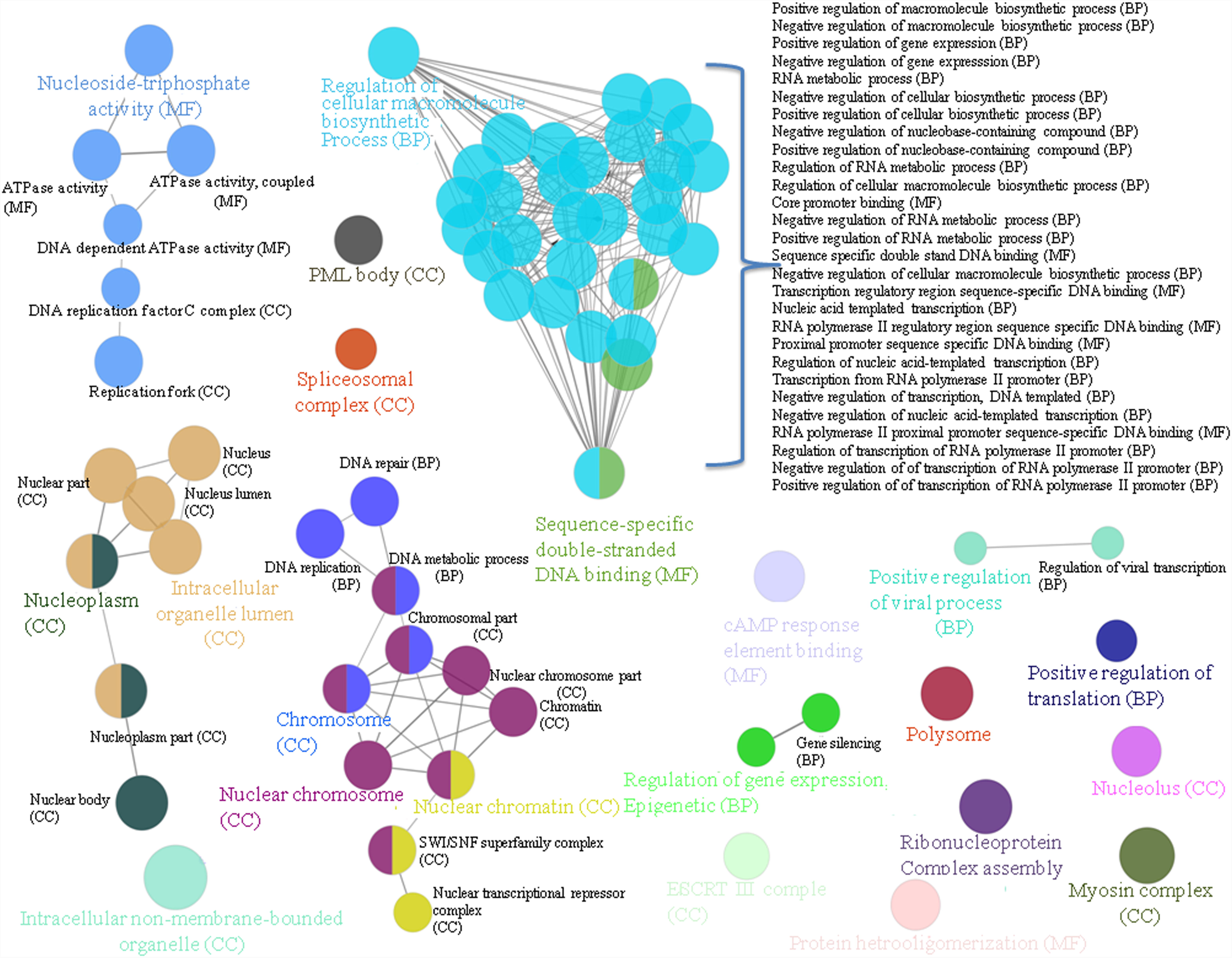
Analysis of statistically significantly gene ontology terms enriched in vGAF protein interactome. Statistically significant GO terms were visualized in a network layout where the nodes corresponds to GO terms and the edges connecting the nodes corresponds to statistically significant proportion of common protein between those terms. Node size corresponds to term p value. GO terms with highest number of protein in a cluster are colored for emphasis. CC, MF and BP corresponds to cellular compartment, molecular function and biological process and are appropriately abbreviated in front of each GO term.

**Table 2.**
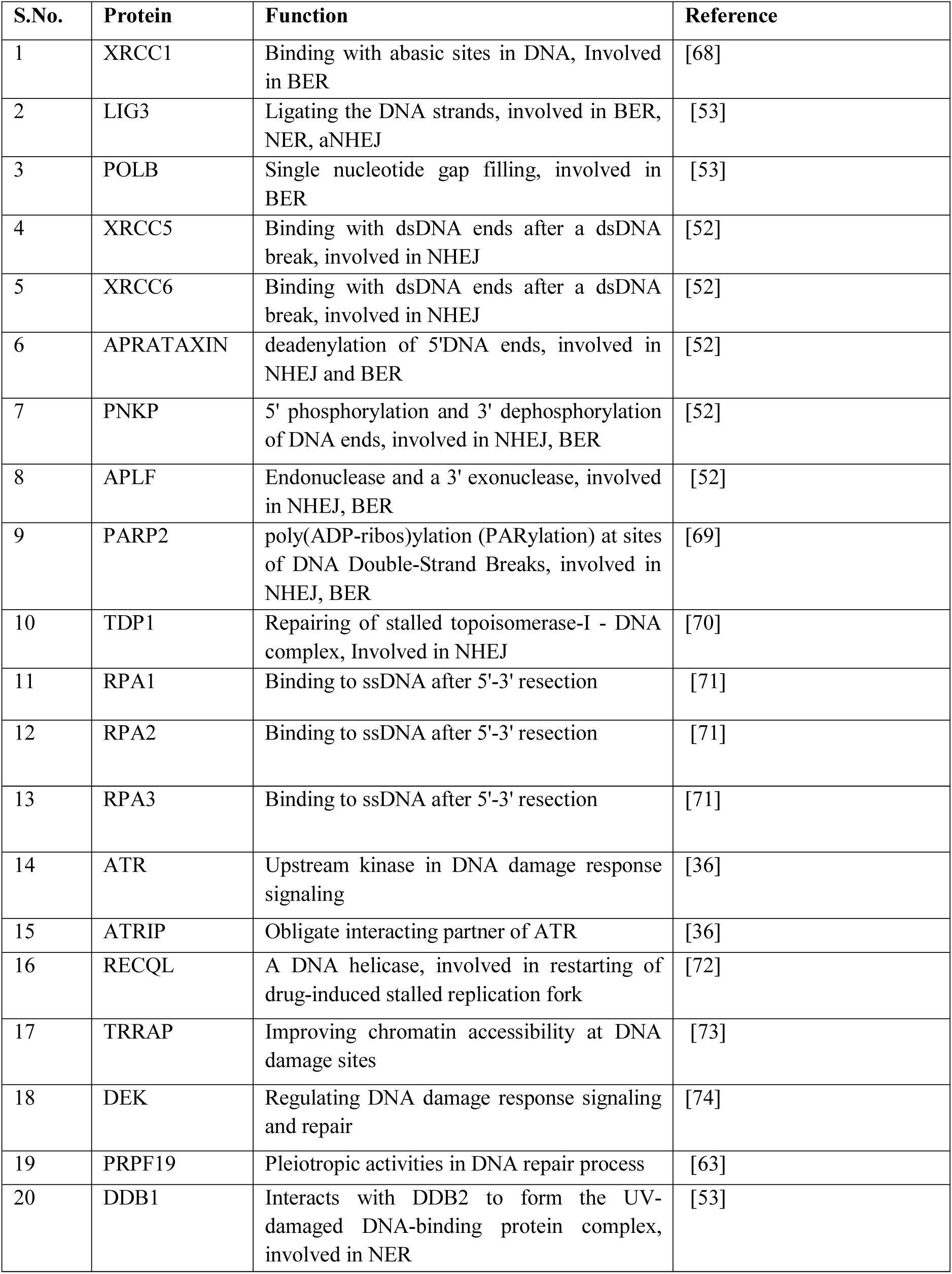
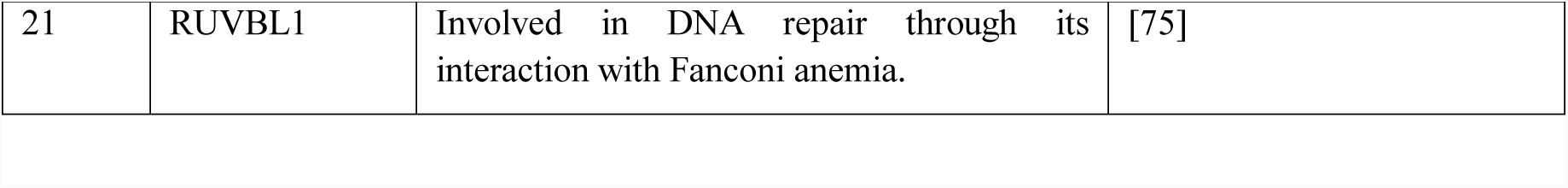
List of DNA repair proteins identified in vGAF protein interactome.

| S.No. | Protein | Function | Reference |
| --- | --- | --- | --- |
| 1 | XRCC1 | Binding with abasic sites in DNA, Involved in BER | [68] |
| 2 | LIG3 | Ligating the DNA strands, involved in BER, NER, aNHEJ | [53] |
| 3 | POLB | Single nucleotide gap filling, involved in BER | [53] |
| 4 | XRCC5 | Binding with dsDNA ends after a dsDNA break, involved in NHEJ | [52] |
| 5 | XRCC6 | Binding with dsDNA ends after a dsDNA break, involved in NHEJ | [52] |
| 6 | APRATAXIN | deadenylation of 5'DNA ends, involved in NHEJ and BER | [52] |
| 7 | PNKP | 5' phosphorylation and 3' dephosphorylation of DNA ends, involved in NHEJ, BER | [52] |
| 8 | APLF | Endonuclease and a 3' exonuclease, involved in NHEJ, BER | [52] |
| 9 | PARP2 | poly(ADP-ribos)ylation (PARylation) at sites of DNA Double-Strand Breaks, involved in NHEJ, BER | [69] |
| 10 | TDP1 | Repairing of stalled topoisomerase-I - DNA complex, Involved in NHEJ | [70] |
| 11 | RPA1 | Binding to ssDNA after 5'-3' resection | [71] |
| 12 | RPA2 | Binding to ssDNA after 5'-3' resection | [71] |
| 13 | RPA3 | Binding to ssDNA after 5'-3' resection | [71] |
| 14 | ATR | Upstream kinase in DNA damage response signaling | [36] |
| 15 | ATRIP | Obligate interacting partner of ATR | [36] |
| 16 | RECQL | A DNA helicase, involved in restarting of drug-induced stalled replication fork | [72] |
| 17 | TRRAP | Improving chromatin accessibility at DNA damage sites | [73] |
| 18 | DEK | Regulating DNA damage response signaling and repair | [74] |
| 19 | PRPF19 | Pleiotropic activities in DNA repair process | [63] |
| 20 | DDB1 | Interacts with DDB2 to form the UV-damaged DNA-binding protein complex, involved in NER | [53] |
| 21 | RUVBL1 | Involved in DNA repair through its interaction with Fanconi anemia. | [75] |

### 3.3 GAF associates with ATP dependent chromatin remodelers

ATP dependent chromatin remodelers (ADCRs) are key players in the regulation of chromatin packaging and thereby essential for various DNA dependent nuclear functions [34]. There are four different families of chromatin remodeling complexes viz. SWI/SNF, ISWI, CHD and INO80 [29]. All these complexes have an ATPase subunit and a number of non-catalytic subunits, which further diversify these complexes into sub-families [29]. In our analysis, we identify all the components of NuRD complex that belongs to the CHD family of ADCRs (Table 1). We identify Mi-2beta/CHD4 that is the ATPase subunit of this complex, along with that, we also identify all other non-catalytic subunits viz. MBD3, HDAC1, HDAC2, RbAp46, p66-alpha/beta and DOC-1. Similarly, we identify both the components of ACF subfamily of ISWI complexes, viz. SNF2H (ATPase subunit) and BAZ1A (non-catalytic subunit) [29]. On the contrary, we do not identify most of the components of SWI/SNF and INO80 complexes, other than BAF53a, ARID1A (SWI/SNF and INO80) & RUVBL1 (INO80) [29]. These results suggest that vGAF interacts with whole ACF and NuRD remodeling complexes.

### 3.4 vGAF interacts with a spectrum of DNA repair proteins and is essential for efficient DNA repair and cell survival after UV induced DNA damage

Cellular response to the DNA damage is mediated by DNA damage response (DDR) signaling pathways. ATM (ataxia-telangiectasia mutated), ATR (ATM- and Rad3-Related), and DNA-PKcs (DNA-dependent protein kinase) are the most upstream kinases of DDR signaling pathway [35]. We identify ATR and its obligate interactor ATRIP in our vGAF IP mass-spec data. ATR is of significant importance as it responds to both double strand DNA breaks and a variety of DNA lesions that interfere with DNA replication [36,37]. Further, we find all three RPA70, RPA14 and RPA32 proteins that form complex with ssDNA and recruit ATR/ATRIP complex to the site of DNA damage/replication fork [38]. We find several other DNA repair proteins that are involved in DNA break repair including Ku60 and Ku80, suggesting a physical association between vGAF and DNA repair machinery (Table 2). The phosphorylated form of histone variant H2AX (γ-H2AX) quickly accumulates at the site of double strand DNA breaks within minutes after the induction of DNA damage [39]. It has extensively been used both as a spatial and quantitative marker for DNA damage [39]. In order to determine whether vGAF co-localizes to the sites of DNA double strand breaks, we performed immunostaining of γ-H2AX 1-hour after the UV treatment of C2C12 myoblast cells expressing FLAG-vGAF. As shown in Fig. 4a, co-immunostaining of γ-H2AX (green) and FLAG-vGAF (magenta) shows several regions where these two proteins co-localize with each other, suggesting an association of vGAF with doubled strand DNA break lesions. To further substantiate our results, we used tissues and cells derived from vGAF knock out mice and asked if vGAF could affect the process of DNA-damage repair. We measured the levels of γ-H2AX using western blotting and used it as a proxy for the level of DNA damage in the tissues or cells used for protein extract preparation. vGAF has been shown to express predominantly in adult mouse skin [12,13]. Protein extracts prepared 3-hours after UV treatment of the skin explants shows a higher accumulation of γ-H2AX in vGAF KO skin compared to the wild-type controls, suggesting a delayed repair of DNA-damage in KO skin tissue compared to wild-type tissue (Fig. 4a). We further checked the survival of skin primary fibroblast 4-days after the UV treatment using MTT assay. Here again, we see that primary fibroblast cells from vGAF KO skin are more sensitive to UV treatment compared to the control (Fig. 4b). To further confirm our results, we derived primary fibroblast cells from the lungs which is another organ where vGAF show a significant expression [40]. Protein extracts prepared 3-hours after UV treatment of primary lung fibroblasts show an increased signal of γ-H2AX in western blots compared to the protein extract from the wild-type lung primary fibroblast cells (Fig. 4c). Collectively, these results show that vGAF is essential for efficient DNA repair after UV induced DNA damage. Furthermore, these results serve as a biological validation of our IP mass-spec data that suggest a physical association between vGAF and DNA repair machinery.

**Fig. 4.**
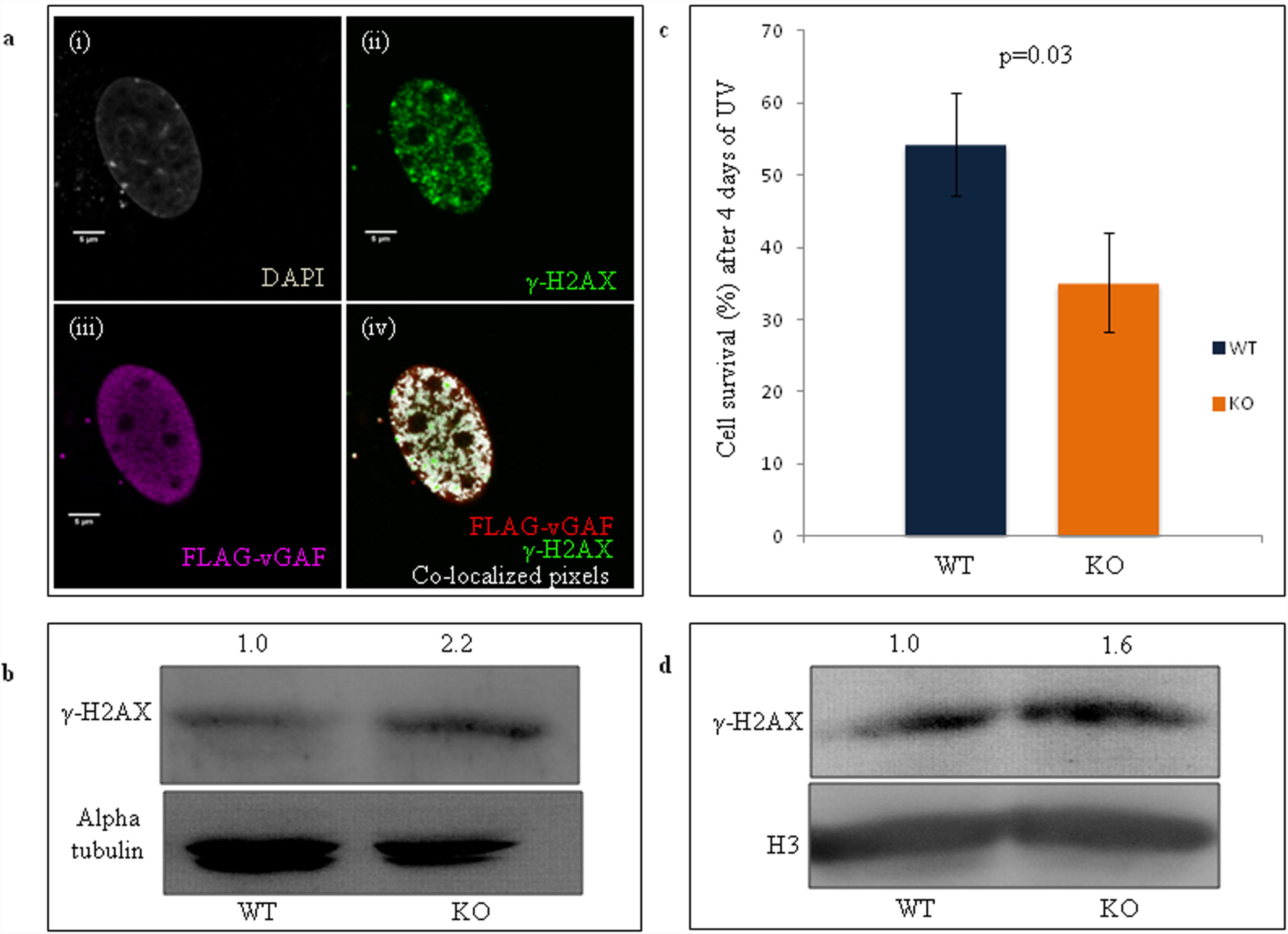
vGAF co-localizes with DNA double strand break lesions sites and it is essential for efficient DNA repair and cell survival after UV induced DNA damage. [a] C2C12 myoblast cells show co-localization of FLAG-vGAF with γ-H2A.X. C2C12 myoblast cells were transfected with FLAG-vGAF and 36 hours post-transfection cells were treated with UVC (5J/m2) and allowed to recover for 1 hour. Cells were co-immunostained with anti-γ-H2A.X and anti-FLAG antibodies. Single optical section from the center of nucleus show the staining of (i) DAPI (grey), (ii) FLAG-vGAF (magenta) (iii) γ-H2A.X (green) and (iv) co-localization of FLAG-vGAF and γ-H2A.X (white pixels). ImageJ was used to identify and highlight pixels where FLAG-vGAF and γ-H2A.X staining signals were present together. Scale bars are 5 micrometer. [b] vGAF KO mice skin explants show higher levels of DNA damage (increased Y-H2Ax) after UV induced DNA damage. Skin explants were treated with UVC (60J/m2) and were allowed to recover for 3 hours. Increase in Y-H2Ax was detected in protein extracts prepared from vGAF KO mice skin explants. [c] Skin primary fibroblast cells derived from vGAF KO mice are sensitive to UVC treatment. Skin primary fibroblast cells were treated with UVC (30J/m2). Cell survival assay shows a significant decrease in survival of primary fibroblast derived from skin of KO mice. [d] Lung primary fibroblasts derived from vGAF KO mice show higher levels of DNA damage (increase in Y-H2Ax) after UV induced DNA damage. Lung primary fibroblast were treated with UVC (5J/m2) and were allowed to recover for 3 hours. Increased Y-H2Ax was detected in protein extracts prepared from cells derived from vGAF KO.

### 3.5 Expression of vGAF is downregulated in skin cancer tissue

UV induced DNA damage is the primary cause of skin cancers and that prompted us to compare the expression data of vGAF (ZBTB7B) and DNA repair proteins that interact with it (Table 2), in skin cutaneous melanoma (SCM) and normal skin tissue (NST). GEPIA (Gene Expression Profiling Interactive Analysis) is a web-based tool for the analysis of TCGA (The Cancer Genome Atlas) and GTEx (Genotype-Tissue Expression) database [41]. We analyzed gene expression data from these two databases using GEPIA and show that SCM tissue samples express vGAF at a much lower level than the NST samples (Fig. 5a), while other DNA repair proteins which were taken in this study are either upregulated or show no significant difference in expression in SCM vs. NST (Sup. Fig. 4). Further analysis also shows that vGAF expression does not change significantly across all the stages of SCM suggesting a consistent downregulation of vGAF even in the stage-0 of SCM (Fig. 5b). Collectively, these results suggest that vGAF is heavily downregulated in SCM even at stage-0, and thus low levels of vGAF in skin biopsies can be used as a diagnostic biomarker for SCM.

**Fig. 5.**
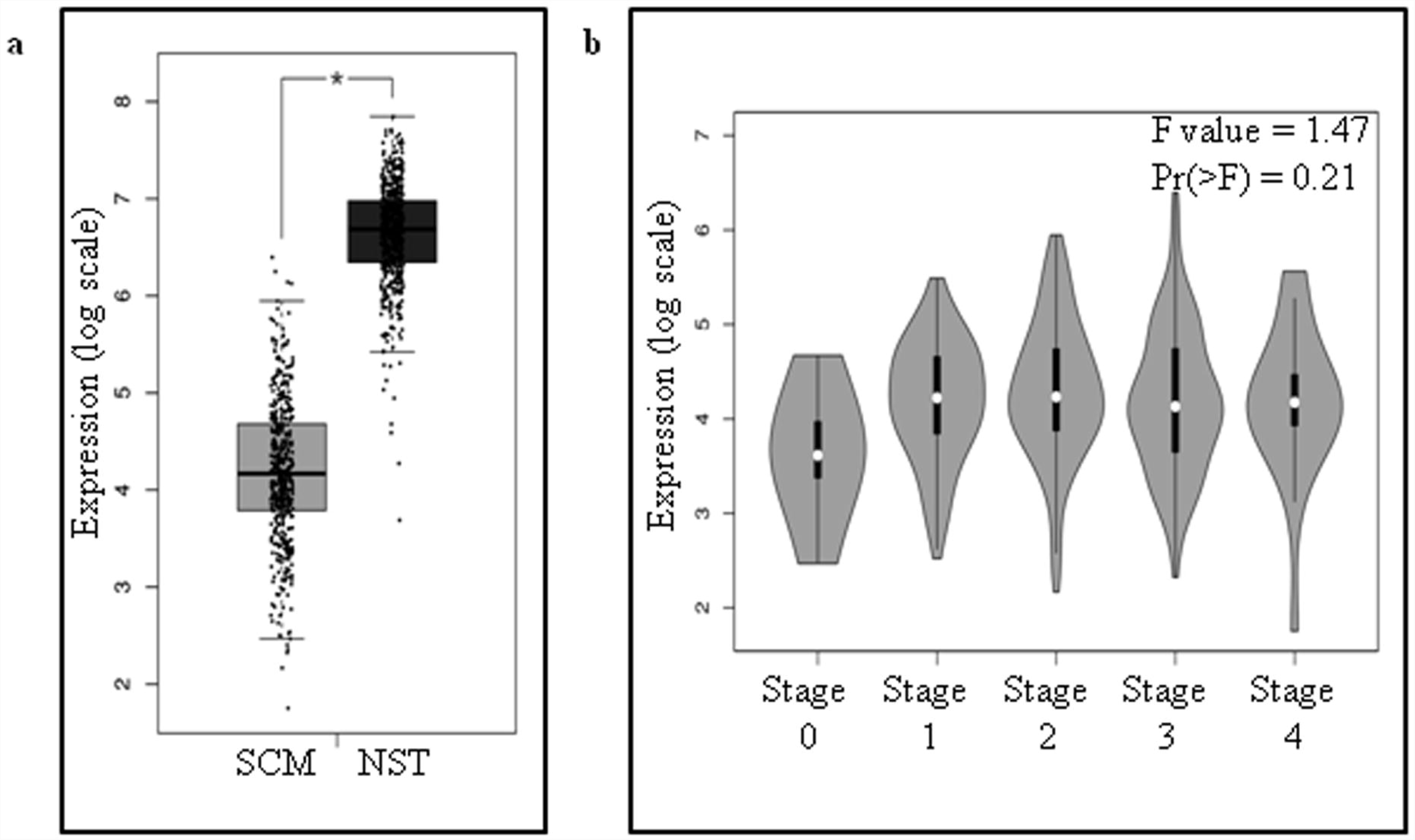
Analysis of vGAF expression in skin cutaneous melanoma (SCM) samples from human subjects. [a] vGAF expression is downregulated in SCM samples compared to normal skin tissue (NST). GEPIA is used to analyze the expression data from 461 SCM and 558 NST samples. Analysis shows a significant decrease in vGAF expression in SCM (p<=0.001). [b] vGAF expression levels does not change significantly across different stages of SCM. Violin plots of vGAF expression data across pathological stages of SCM shows a consistent downregulation including stage 0 of SCM.

## 4. Discussion

Gene-knockout mouse model and extensive cell culture studies have revealed functions of vGAF in a variety of biological processes viz. development of CD4+ T-cells, brown-fat development, lactation and thermogenesis [8,9,10,11]. Intriguingly, the molecular functions of vGAF that form the basis for these biological processes are confounding in nature. Studies suggest that vGAF is a repressor of *cd8* gene in CD4+ T-cells while vGAF forms a ribo-nucleoprotein complex with lincRNA to activate thermogenic gene expression program in adipocytes [16,10]. Moreover, several cell culture based reports on vGAF indicate an even a broader array of molecular activities associated with vGAF like nuclear lamina mediated repression of key developmental genes and enhancer-blocking activity in mammalian *Hox* clusters [19,20]. Studies in both *Drosophila melanogaster* and mammals have suggested the importance of protein-protein interactions in the molecular processes associated with GAF proteins [16,20,22-24,42-44]. Here, we have identified a comprehensive protein interactome of vGAF to understand the diversity of vGAF protein interacting complexes.

Existing literature on vGAF shows that it binds on the promoter of genes like *UDP glucose dehydrogenase, eomesodermin, Col1a1, Socs1 and Cish* and regulates the gene expression [15,17,18,44]. In our study, we also identify a number of transcription factors and components of RNA-polymerase macromolecular complexes. The interaction of vGAF with these protein complexes can modulate the expression of genes where it binds on promoters. Intriguingly, we identify several proteins that are involved in RNA metabolic processes like alternative splicing & mRNA maturation. The functional significance of this association between vGAF and components of AS is exemplified by a high-throughput molecular screen which identifies vGAF as one of the factors that affects developmentally important AS events [45]. Gene ontology analysis of vGAF protein interactome also suggests the interaction of vGAF with proteins involved in the regulation of viral life cycle. A recent report shows that knockdown of vGAF in A549 cells results in almost 50% reduction of Influenza A virus (IAV) titers 24 hours post infection (hpi) compared to the mock treatment, suggesting the functional relevance of the proteins identified in our analysis [46].

Unexpectedly, we find components of ESCRT-III complex in our vGAF protein interactome list. ESCRT complex proteins are required for multivesicular body/endosome biogenesis [47]. The specific role of ESCRT-III complex members is in membrane remodeling (budding and scission) reactions during multivesicular body biogenesis [47,48]. Interestingly, recent studies have shown that ESCRT-III complex is implicated in reformation of nuclear envelope, and co-localizes with Lamin-B receptor and LAP2-beta during the assembly of nuclear envelop after mitotic exit [49]. LAP2-beta is a known interactor of vGAF and has been identified in our studies as well [20]. Microscopic studies have also revealed that vGAF binds on the mitotic chromosome and interacts with lamin B1 during the assemble of nuclear envelop [20]. These independent studies further support a physical link between vGAF and ESCRT-III complex.

Our analysis shows an association of vGAF with whole ACF and NuRD chromatin remodeling complexes. Apart from their gene regulatory functions, both ACF and NuRD complexes are involved in DNA replication and repair [50,51]. In our analysis, the DNA repair associated functions of these chromatin remodeling complexes are of vital importance as we also identify a number of other DNA repair proteins that interact with vGAF. We identify both Ku60 and Ku80, the two major proteins that form complex with DNA ends after a double strand DNA break. Furthermore, our analysis shows the presence of several cNHEJ (classical Non-homologous end joining) accessory proteins (PNKP, Aprataxin, APLF) and ligase3, a component of alternative NHEJ [52]. We also identify the components of base excision DNA repair, viz., XRCC1, Polb and ligase3 [53]. These results show a physical association of vGAF with components of DNA repair machinery suggesting a possible functional role of vGAF in DNA repair process. The functional aspects of this physical association between vGAF and DNA repair machinery is further supported by our results that show that vGAF is necessary for efficient DNA repair and cell survival after DNA damage.

The novel role of vGAF in DNA repair process is quite surprising considering the established functions of this protein as a zinc-finger transcription factor with sequence-specific DNA binding ability. However, we have recently started understanding the role of sequence-specific zinc-finger transcription factors in the maintenance of genome integrity [54]. ZBTB7A/LRF, a paralogue of vGAF in mammals, has recently been shown to be involved in cNHEJ DNA repair pathway [55]. Interestingly, LRF is a very well reported interacting partner of vGAF and it is present in our vGAF protein interaction data as well. DNA damage and repair processes has closely been linked to tumorigenesis [56]. UV-induced DNA damage is the major cause of tumor formation in skin tissue where vGAF also expresses predominantly [57]. Our results show that vGAF is heavily downregulated in SCM and its expression does not change significantly across major stages of melanoma progression, suggesting downregulation even in stage 0 of SCM. We speculate that low expression level of vGAF in the cells interferes with the assembly of DNA-repair protein complexes and thus makes the cell less efficient in repairing the DNA lesions.

Recent studies have established that DNA repair is closely linked with two other major nuclear processes viz. transcription and alternative splicing [58,59]. Noticeably, our vGAF protein interactome data includes several proteins, which are uniquely or redundantly involved in these three processes. For example, vGAF interacts with factors (like RBM14, RBMX and PRPF19) that are involved in both splicing/RNA processing and DNA damage repair [30,31,60-63]. Likewise, our analysis shows that vGAF associates with transcription factors which are involved in DNA damage repair (LRF, ZNF281) and RNA processing (LRF, YY1) [45,55,64-67]. Altogether, these observations suggest that vGAF can influence all three major nuclear processes (Transcription, Splicing and DNA repair) and stands a central position in nuclear-proteome-wide interaction network where molecular circuits of transcription, splicing and DNA repair criss-cross each other.

## 5. Conclusion

We report a comprehensive protein interactome of vGAF in C2C12 myoblast cells. Analysis of the interactome suggests that vGAF interacts with a functionally diverse set of proteins belonging to transcriptional regulation, chromatin remodeling, RNA metabolism, viral life cycle and DNA repair. We unravel an important function of vGAF in DNA repair and cell survival after UV induced DNA damage. Our study shows that vGAF is downregulated in all the major stage of SCM and, therefore, can be used as an early diagnostic biomarker. Taken together, our study provides a molecular basis for the diverse function of vGAF and also indicates its unexplored functions in biological processes.

## 6. Acknowledgment

AS acknowledge GATE-SRF fellowship from CSIR towards the PhD program. Authors thank CCMB-Proteomics facility members for performing the mass-spectrometry.

## Abbreviations

ACF: ATP-utilizing chromatin assembly and remodeling factor)
ADCR: ATP dependent chromatin remodelers
AS: Alternative Splicing
BER: Base excision repair
BTB/POZ: Broad-Complex, Tramtrack and Bric a brac/POxvirus and Zinc finger domain
DmGAF: *Drosophila melanogaster* GAF
GAF: GAGA associated Factor
GO: Gene ontology
GOBP: Gene Ontology biological process
GOCC: Gene ontology cellular compartment
GOMF: Gene ontology molecular function
ISWI: Imitation SWI
lincRNA: Long intergenic non-coding RNA
MTT: 3-(4,5-dimethylthiazol-2-yl)-2,5-diphenyltetrazolium bromide
NER: Nucleotide excision repair
NHEJ: Non-homologous end joining
NST: Normal skin tissue
NuRD: Nucleosome remodeling and deacetylase
SCM: Skin cutaneous melanoma
SWI/SNF: SWItch/Sucrose Non-Fermentable
ThPOK: T helper POK
vGAF: Vertebrate GAF

## References

1. Farkas G, Gausz J, Galloni M, Reuter G, Gyurkovics H, Karch F (1994) The Trithorax-like gene encodes the Drosophila GAGA factor. Nature 371 (6500):806–808. doi:10.1038/371806a0

2. Bhat KM, Farkas G, Karch F, Gyurkovics H, Gausz J, Schedl P (1996) The GAGA factor is required in the early Drosophila embryo not only for transcriptional regulation but also for nuclear division. Development 122 (4):1113–1124

3. Biggin MD, Tjian R (1988) Transcription factors that activate the Ultrabithorax promoter in developmentally staged extracts. Cell 53 (5):699–711

4. Lu Q, Wallrath LL, Granok H, Elgin SC (1993) (CT)n (GA)n repeats and heat shock elements have distinct roles in chromatin structure and transcriptional activation of the Drosophila hsp26 gene. Mol Cell Biol 13 (5):2802–2814

5. Mishra RK, Mihaly J, Barges S, Spierer A, Karch F, Hagstrom K, Schweinsberg SE, Schedl P (2001) The iab-7 polycomb response element maps to a nucleosome-free region of chromatin and requires both GAGA and pleiohomeotic for silencing activity. Mol Cell Biol 21 (4):1311–1318. doi:10.1128/MCB.21.4.1311-1318.2001

6. Schweinsberg S, Hagstrom K, Gohl D, Schedl P, Kumar RP, Mishra R, Karch F (2004) The enhancer-blocking activity of the Fab-7 boundary from the Drosophila bithorax complex requires GAGA-factor-binding sites. Genetics 168 (3):1371–1384. doi:10.1534/genetics.104.029561

7. Matharu NK, Hussain T, Sankaranarayanan R, Mishra RK (2010) Vertebrate homologue of Drosophila GAGA factor. J Mol Biol 400 (3):434–447. doi:10.1016/j.jmb.2010.05.010

8. Egawa T, Littman DR (2008) ThPOK acts late in specification of the helper T cell lineage and suppresses Runx-mediated commitment to the cytotoxic T cell lineage. Nat Immunol 9 (10):1131–1139. doi:10.1038/ni.1652

9. Muroi S, Naoe Y, Miyamoto C, Akiyama K, Ikawa T, Masuda K, Kawamoto H, Taniuchi I (2008) Cascading suppression of transcriptional silencers by ThPOK seals helper T cell fate. Nat Immunol 9 (10):1113–1121. doi:10.1038/ni.1650

10. Li S, Mi L, Yu L, Yu Q, Liu T, Wang GX, Zhao XY, Wu J, Lin JD (2017) Zbtb7b engages the long noncoding RNA Blnc1 to drive brown and beige fat development and thermogenesis. Proc Natl Acad Sci U S A 114 (34):E7111–E7120. doi:10.1073/pnas.1703494114

11. Zhang R, Ma H, Gao Y, Wu Y, Qiao Y, Geng A, Cai C, Han Y, Zeng YA, Liu X, Ge G (2018) Th-POK regulates mammary gland lactation through mTOR-SREBP pathway. PLoS Genet 14 (2):e1007211. doi:10.1371/journal.pgen.1007211

12. Galera P, Musso M, Ducy P, Karsenty G (1994) c-Krox, a transcriptional regulator of type I collagen gene expression, is preferentially expressed in skin. Proc Natl Acad Sci U S A 91 (20):9372–9376

13. Galera P, Park RW, Ducy P, Mattei MG, Karsenty G (1996) c-Krox binds to several sites in the promoter of both mouse type I collagen genes. Structure/function study and developmental expression analysis. J Biol Chem 271 (35):21331–21339

14. Stratigi K, Kapsetaki M, Aivaliotis M, Town T, Flavell RA, Spilianakis CG (2015) Spatial proximity of homologous alleles and long noncoding RNAs regulate a switch in allelic gene expression. Proc Natl Acad Sci U S A 112 (13):E1577–1586. doi:10.1073/pnas.1502182112

15. Luckey MA, Kimura MY, Waickman AT, Feigenbaum L, Singer A, Park JH (2014) The transcription factor ThPOK suppresses Runx3 and imposes CD4(+) lineage fate by inducing the SOCS suppressors of cytokine signaling. Nat Immunol 15 (7):638–645. doi:10.1038/ni.2917

16. Rui J, Liu H, Zhu X, Cui Y, Liu X (2012) Epigenetic silencing of CD8 genes by ThPOK-mediated deacetylation during CD4 T cell differentiation. J Immunol 189 (3):1380–1390. doi:10.4049/jimmunol.1201077

17. Li Y, Tsun A, Gao Z, Han Z, Gao Y, Li Z, Lin F, Wang Y, Wei G, Yao Z, Li B (2013) 60-kDa Tat-interactive protein (TIP60) positively regulates Th-inducing POK (ThPOK)-mediated repression of eomesodermin in human CD4+ T cells. J Biol Chem 288 (22):15537–15546. doi:10.1074/jbc.M112.430207

18. Beauchef G, Kypriotou M, Chadjichristos C, Widom RL, Poree B, Renard E, Moslemi S, Wegrowski Y, Maquart FX, Pujol JP, Galera P (2005) c-Krox down-regulates the expression of UDP-glucose dehydrogenase in chondrocytes. Biochem Biophys Res Commun 333 (4):1123–1131. doi:10.1016/j.bbrc.2005.06.020

19. Srivastava S, Puri D, Garapati HS, Dhawan J, Mishra RK (2013) Vertebrate GAGA factor associated insulator elements demarcate homeotic genes in the HOX clusters. Epigenetics Chromatin 6 (1):8. doi:10.1186/1756-8935-6-8

20. Zullo JM, Demarco IA, Pique-Regi R, Gaffney DJ, Epstein CB, Spooner CJ, Luperchio TR, Bernstein BE, Pritchard JK, Reddy KL, Singh H (2012) DNA sequence-dependent compartmentalization and silencing of chromatin at the nuclear lamina. Cell 149 (7):1474–1487. doi:10.1016/j.cell.2012.04.035

21. Srivastava A, Kumar AS, Mishra RK (2018) Vertebrate GAF/ThPOK: emerging functions in chromatin architecture and transcriptional regulation. Cell Mol Life Sci 75 (4):623–633. doi:10.1007/s00018-017-2633-7

22. Lomaev D, Mikhailova A, Erokhin M, Shaposhnikov AV, Moresco JJ, Blokhina T, Wolle D, Aoki T, Ryabykh V, Yates JR, 3rd, Shidlovskii YV, Georgiev P, Schedl P, Chetverina D (2017) The GAGA factor regulatory network: Identification of GAGA factor associated proteins. PLoS One 12 (3):e0173602. doi:10.1371/journal.pone.0173602

23. Chopra VS, Srinivasan A, Kumar RP, Mishra K, Basquin D, Docquier M, Seum C, Pauli D, Mishra RK (2008) Transcriptional activation by GAGA factor is through its direct interaction with dmTAF3. Dev Biol 317 (2):660–670. doi:10.1016/j.ydbio.2008.02.008

24. Mishra K, Chopra VS, Srinivasan A, Mishra RK (2003) Trl-GAGA directly interacts with lola like and both are part of the repressive complex of Polycomb group of genes. Mech Dev 120 (6):681–689

25. Seluanov A, Vaidya A, Gorbunova V (2010) Establishing primary adult fibroblast cultures from rodents. J Vis Exp (44). doi:10.3791/2033

26. Kallappagoudar S, Varma P, Pathak RU, Senthilkumar R, Mishra RK (2010) Nuclear Matrix Proteome Analysis of Drosophila melanogaster. Mol Cell Proteomics 9 (9):2005–2018. doi:10.1074/mcp.M110.001362

27. Bindea G, Mlecnik B, Hackl H, Charoentong P, Tosolini M, Kirilovsky A, Fridman WH, Pages F, Trajanoski Z, Galon J (2009) ClueGO: a Cytoscape plug-in to decipher functionally grouped gene ontology and pathway annotation networks. Bioinformatics 25 (8):1091–1093. doi:10.1093/bioinformatics/btp101

28. Shannon P, Markiel A, Ozier O, Baliga NS, Wang JT, Ramage D, Amin N, Schwikowski B, Ideker T (2003) Cytoscape: a software environment for integrated models of biomolecular interaction networks. Genome Res 13 (11):2498–2504. doi:10.1101/gr.1239303

29. Clapier CR, Cairns BR (2009) The biology of chromatin remodeling complexes. Annu Rev Biochem 78:273–304. doi:10.1146/annurev.biochem.77.062706.153223

30. Simon NE, Yuan M, Kai M (2017) RNA-binding protein RBM14 regulates dissociation and association of non-homologous end joining proteins. Cell Cycle 16 (12):1175–1180. doi:10.1080/15384101.2017.1317419

31. Auboeuf D, Dowhan DH, Li X, Larkin K, Ko L, Berget SM, O’Malley BW (2004) CoAA, a nuclear receptor coactivator protein at the interface of transcriptional coactivation and RNA splicing. Mol Cell Biol 24 (1):442–453

32. Maeng YS, Kwon JY, Kim EK, Kwon YG (2015) Heterochromatin Protein 1 Alpha (HP1alpha: CBX5) is a Key Regulator in Differentiation of Endothelial Progenitor Cells to Endothelial Cells. Stem Cells 33 (5):1512–1522. doi:10.1002/stem.1954

33. Oeckinghaus A, Ghosh S (2009) The NF-kappaB family of transcription factors and its regulation. Cold Spring Harb Perspect Biol 1 (4):a000034. doi:10.1101/cshperspect.a000034

34. Bartholomew B (2014) Regulating the chromatin landscape: structural and mechanistic perspectives. Annu Rev Biochem 83:671–696. doi:10.1146/annurev-biochem-051810-093157

35. Blackford AN, Jackson SP (2017) ATM, ATR, and DNA-PK: The Trinity at the Heart of the DNA Damage Response. Mol Cell 66 (6):801–817. doi:10.1016/j.molcel.2017.05.015

36. Cortez D, Guntuku S, Qin J, Elledge SJ (2001) ATR and ATRIP: partners in checkpoint signaling. Science 294 (5547):1713–1716. doi:10.1126/science.1065521

37. Flynn RL, Zou L (2011) ATR: a master conductor of cellular responses to DNA replication stress. Trends Biochem Sci 36 (3):133–140. doi:10.1016/j.tibs.2010.09.005

38. Zou L, Elledge SJ (2003) Sensing DNA damage through ATRIP recognition of RPA-ssDNA complexes. Science 300 (5625):1542–1548. doi:10.1126/science.1083430

39. Kuo LJ, Yang LX (2008) Gamma-H2AX - a novel biomarker for DNA double-strand breaks. In Vivo 22 (3):305–309

40. Michaloski JS, Galante PA, Nagai MH, Armelin-Correa L, Chien MS, Matsunami H, Malnic B (2011) Common promoter elements in odorant and vomeronasal receptor genes. PLoS One 6 (12):e29065. doi:10.1371/journal.pone.0029065

41. Tang Z, Li C, Kang B, Gao G, Li C, Zhang Z (2017) GEPIA: a web server for cancer and normal gene expression profiling and interactive analyses. Nucleic Acids Res 45 (W1):W98–W102. doi:10.1093/nar/gkx247

42. Widom RL, Lee JY, Joseph C, Gordon-Froome I, Korn JH (2001) The hcKrox gene family regulates multiple extracellular matrix genes. Matrix Biol 20 (7):451–462

43. Beauchef G, Bigot N, Kypriotou M, Renard E, Poree B, Widom R, Dompmartin-Blanchere A, Oddos T, Maquart FX, Demoor M, Boumediene K, Galera P (2012) The p65 subunit of NF-kappaB inhibits COL1A1 gene transcription in human dermal and scleroderma fibroblasts through its recruitment on promoter by protein interaction with transcriptional activators (c-Krox, Sp1, and Sp3). J Biol Chem 287 (5):3462–3478. doi:10.1074/jbc.M111.286443

44. Kypriotou M, Beauchef G, Chadjichristos C, Widom R, Renard E, Jimenez SA, Korn J, Maquart FX, Oddos T, Von Stetten O, Pujol JP, Galera P (2007) Human collagen Krox up-regulates type I collagen expression in normal and scleroderma fibroblasts through interaction with Sp1 and Sp3 transcription factors. J Biol Chem 282 (44):32000–32014. doi:10.1074/jbc.M705197200

45. Han H, Braunschweig U, Gonatopoulos-Pournatzis T, Weatheritt RJ, Hirsch CL, Ha KCH, Radovani E, Nabeel-Shah S, Sterne-Weiler T, Wang J, O’Hanlon D, Pan Q, Ray D, Zheng H, Vizeacoumar F, Datti A, Magomedova L, Cummins CL, Hughes TR, Greenblatt JF, Wrana JL, Moffat J, Blencowe BJ (2017) Multilayered Control of Alternative Splicing Regulatory Networks by Transcription Factors. Mol Cell 65 (3):539–553 e537. doi:10.1016/j.molcel.2017.01.011

46. Chen SC, Jeng KS, Lai MMC (2017) Zinc Finger-Containing Cellular Transcription Corepressor ZBTB25 Promotes Influenza Virus RNA Transcription and Is a Target for Zinc Ejector Drugs. J Virol 91 (20). doi:10.1128/JVI.00842-17

47. McCullough J, Colf LA, Sundquist WI (2013) Membrane fission reactions of the mammalian ESCRT pathway. Annu Rev Biochem 82:663–692. doi:10.1146/annurev-biochem-072909-101058

48. Raab M, Gentili M, de Belly H, Thiam HR, Vargas P, Jimenez AJ, Lautenschlaeger F, Voituriez R, Lennon-Dumenil AM, Manel N, Piel M (2016) ESCRT III repairs nuclear envelope ruptures during cell migration to limit DNA damage and cell death. Science 352 (6283):359–362. doi:10.1126/science.aad7611

49. Olmos Y, Hodgson L, Mantell J, Verkade P, Carlton JG (2015) ESCRT-III controls nuclear envelope reformation. Nature 522 (7555):236–239. doi:10.1038/nature14503

50. Aydin OZ, Vermeulen W, Lans H (2014) ISWI chromatin remodeling complexes in the DNA damage response. Cell Cycle 13 (19):3016–3025. doi:10.4161/15384101.2014.956551

51. Smeenk G, Wiegant WW, Vrolijk H, Solari AP, Pastink A, van Attikum H (2010) The NuRD chromatin-remodeling complex regulates signaling and repair of DNA damage. J Cell Biol 190 (5):741–749. doi:10.1083/jcb.201001048

52. Lieber MR (2010) The mechanism of double-strand DNA break repair by the nonhomologous DNA end-joining pathway. Annu Rev Biochem 79:181–211. doi:10.1146/annurev.biochem.052308.093131

53. Giglia-Mari G, Zotter A, Vermeulen W (2011) DNA damage response. Cold Spring Harb Perspect Biol 3 (1):a000745. doi:10.1101/cshperspect.a000745

54. Vilas CK, Emery LE, Denchi EL, Miller KM (2018) Caught with One’s Zinc Fingers in the Genome Integrity Cookie Jar. Trends Genet 34 (4):313–325. doi:10.1016/j.tig.2017.12.011

55. Liu XS, Chandramouly G, Rass E, Guan Y, Wang G, Hobbs RM, Rajendran A, Xie A, Shah JV, Davis AJ, Scully R, Lunardi A, Pandolfi PP (2015) LRF maintains genome integrity by regulating the non-homologous end joining pathway of DNA repair. Nat Commun 6:8325. doi:10.1038/ncomms9325

56. Halazonetis TD, Gorgoulis VG, Bartek J (2008) An oncogene-induced DNA damage model for cancer development. Science 319 (5868):1352–1355. doi:10.1126/science.1140735

57. Armstrong BK, Kricker A (2001) The epidemiology of UV induced skin cancer. J Photochem Photobiol B 63 (1-3):8–18

58. Naro C, Bielli P, Pagliarini V, Sette C (2015) The interplay between DNA damage response and RNA processing: the unexpected role of splicing factors as gatekeepers of genome stability. Front Genet 6:142. doi:10.3389/fgene.2015.00142

59. Hanawalt PC, Spivak G (2008) Transcription-coupled DNA repair: two decades of progress and surprises. Nat Rev Mol Cell Biol 9 (12):958–970. doi:10.1038/nrm2549

60. Adamson B, Smogorzewska A, Sigoillot FD, King RW, Elledge SJ (2012) A genome-wide homologous recombination screen identifies the RNA-binding protein RBMX as a component of the DNA-damage response. Nat Cell Biol 14 (3):318–328. doi:10.1038/ncb2426

61. Heinrich B, Zhang Z, Raitskin O, Hiller M, Benderska N, Hartmann AM, Bracco L, Elliott D, Ben-Ari S, Soreq H, Sperling J, Sperling R, Stamm S (2009) Heterogeneous nuclear ribonucleoprotein G regulates splice site selection by binding to CC(A/C)-rich regions in pre-mRNA. J Biol Chem 284 (21):14303–14315. doi:10.1074/jbc.M901026200

62. Song EJ, Werner SL, Neubauer J, Stegmeier F, Aspden J, Rio D, Harper JW, Elledge SJ, Kirschner MW, Rape M (2010) The Prp19 complex and the Usp4Sart3 deubiquitinating enzyme control reversible ubiquitination at the spliceosome. Genes Dev 24 (13):1434–1447. doi:10.1101/gad.1925010

63. Mahajan K (2016) hPso4/hPrp19: a critical component of DNA repair and DNA damage checkpoint complexes. Oncogene 35 (18):2279–2286. doi:10.1038/onc.2015.321

64. Wai DC, Shihab M, Low JK, Mackay JP (2016) The zinc fingers of YY1 bind single-stranded RNA with low sequence specificity. Nucleic Acids Res 44 (19):9153–9165. doi:10.1093/nar/gkw590

65. Sigova AA, Abraham BJ, Ji X, Molinie B, Hannett NM, Guo YE, Jangi M, Giallourakis CC, Sharp PA, Young RA (2015) Transcription factor trapping by RNA in gene regulatory elements. Science 350 (6263):978–981. doi:10.1126/science.aad3346

66. Bielli P, Busa R, Di Stasi SM, Munoz MJ, Botti F, Kornblihtt AR, Sette C (2014) The transcription factor FBI-1 inhibits SAM68-mediated BCL-X alternative splicing and apoptosis. EMBO Rep 15 (4):419–427. doi:10.1002/embr.201338241

67. Pieraccioli M, Nicolai S, Antonov A, Somers J, Malewicz M, Melino G, Raschella G (2016) ZNF281 contributes to the DNA damage response by controlling the expression of XRCC2 and XRCC4. Oncogene 35 (20):2592–2601. doi:10.1038/onc.2015.320

68. Nazarkina ZK, Khodyreva SN, Marsin S, Lavrik OI, Radicella JP (2007) XRCC1 interactions with base excision repair DNA intermediates. DNA Repair (Amst) 6 (2):254–264. doi:10.1016/j.dnarep.2006.10.002

69. Ronson GE, Piberger AL, Higgs MR, Olsen AL, Stewart GS, McHugh PJ, Petermann E, Lakin ND (2018) PARP1 and PARP2 stabilise replication forks at base excision repair intermediates through Fbh1-dependent Rad51 regulation. Nat Commun 9 (1):746. doi:10.1038/s41467-018-03159-2

70. Li J, Summerlin M, Nitiss KC, Nitiss JL, Hanakahi LA (2017) TDP1 is required for efficient non-homologous end joining in human cells. DNA Repair (Amst) 60:40–49. doi:10.1016/j.dnarep.2017.10.003

71. Symington LS, Gautier J (2011) Double-strand break end resection and repair pathway choice. Annu Rev Genet 45:247–271. doi:10.1146/annurev-genet-110410-132435

72. Banerjee T, Brosh RM, Jr. (2015) RECQL: a new breast cancer susceptibility gene. Cell Cycle 14 (22):3540–3543. doi:10.1080/15384101.2015.1066539

73. Murr R, Loizou JI, Yang YG, Cuenin C, Li H, Wang ZQ, Herceg Z (2006) Histone acetylation by Trrap-Tip60 modulates loading of repair proteins and repair of DNA double-strand breaks. Nat Cell Biol 8 (1):91–99. doi:10.1038/ncb1343

74. Kavanaugh GM, Wise-Draper TM, Morreale RJ, Morrison MA, Gole B, Schwemberger S, Tichy ED, Lu L, Babcock GF, Wells JM, Drissi R, Bissler JJ, Stambrook PJ, Andreassen PR, Wiesmuller L, Wells SI (2011) The human DEK oncogene regulates DNA damage response signaling and repair. Nucleic Acids Res 39 (17):7465–7476. doi:10.1093/nar/gkr454

75. Rajendra E, Garaycoechea JI, Patel KJ, Passmore LA (2014) Abundance of the Fanconi anaemia core complex is regulated by the RuvBL1 and RuvBL2 AAA+ ATPases. Nucleic Acids Res 42 (22):13736–13748. doi:10.1093/nar/gku1230

