## Supplementary material for "Interactome of vertebrate GAF/ThPOK reveals its diverse functions in gene regulation and DNA repair": Sup. Fig.

**Supplementary Figure 1**

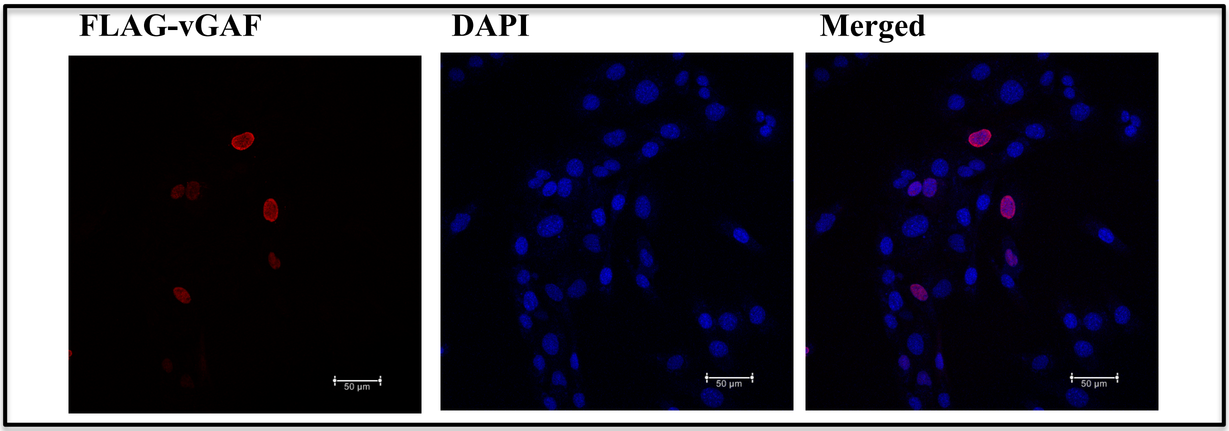


**Supplementary Fig. 1 Nuclear localization of FLAG-vGAF protein in transfected cells**
C2C12 cells transfected with FLAG-vGAF expression plasmids were immunostained with FLAG antibody to detect the localization of FLAG-vGAF. Merged image shows an overlap of FLAG (cy3) and DAPI signals. (Scale bar 50um).

**Supplementary Figure 2**

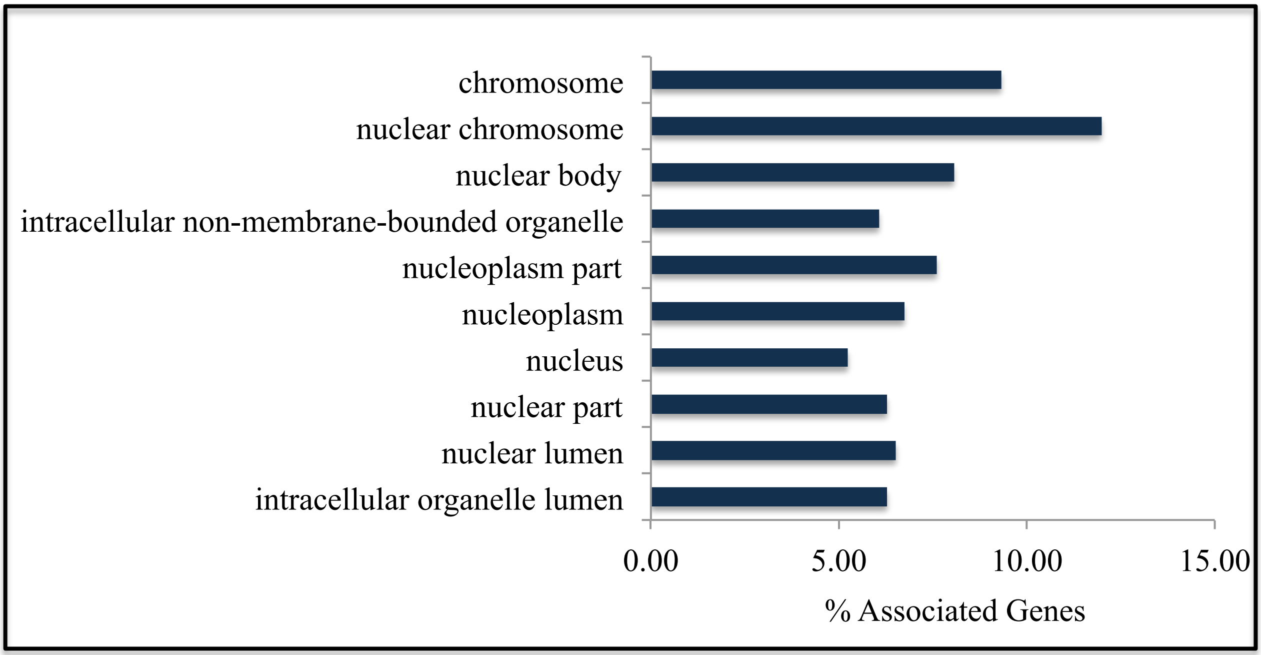


**Supplementary Fig. 2 GO enrichment analysis of immuno-precipitated proteins**
Enrichment test for vGAF protein interactome using ClueGO (see supplementary table 2). Ten most significant gene ontology (cellular component) terms enriched in vGAF protein interactome data are plotted. Y-axis represents the percentage of genes associated with each GO term.

**Supplementary Figure 3**

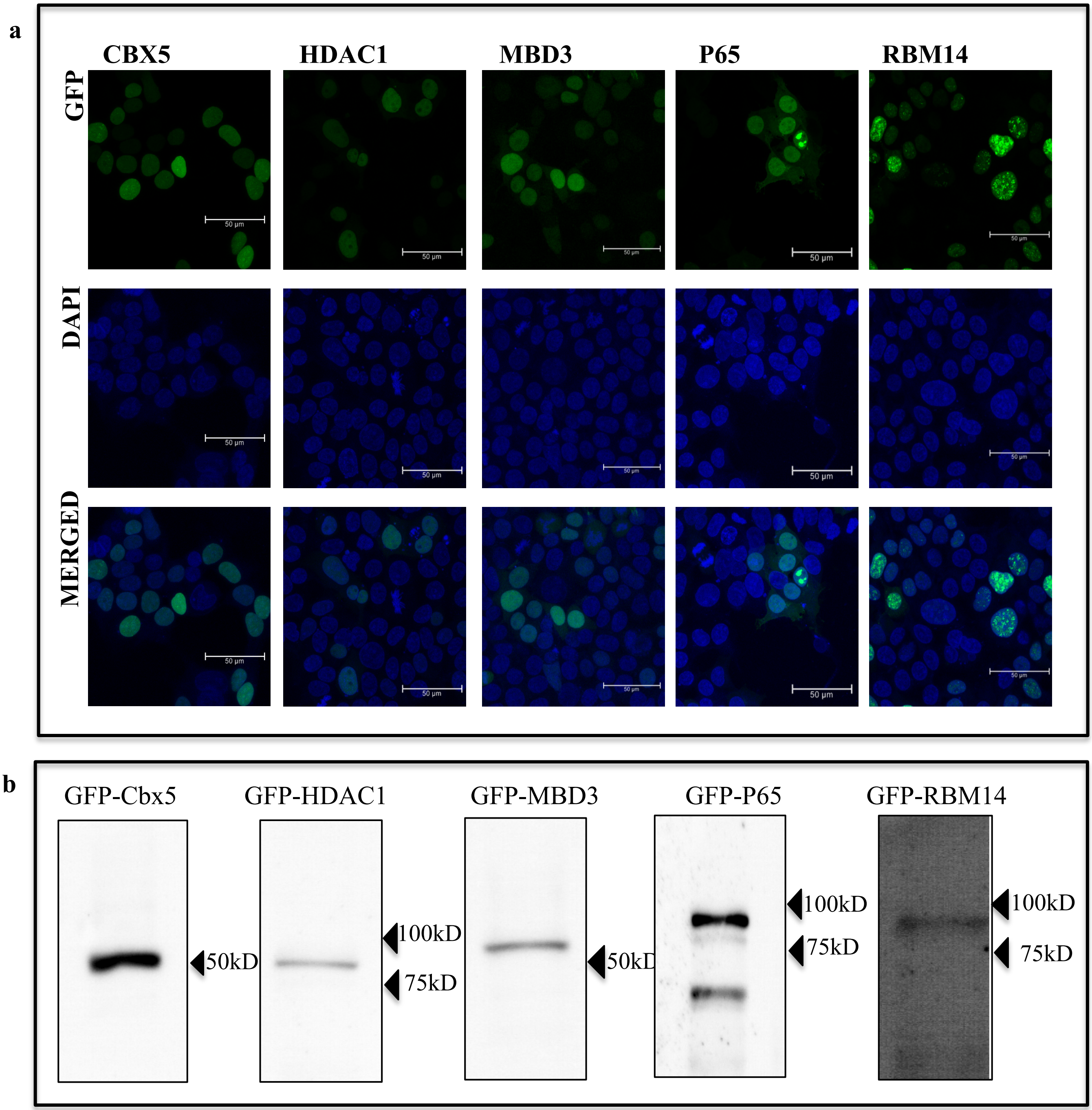


**Supplementary Fig. 3 Expression of N-terminal GFP tagged constructs of candidate proteins (Cbx5, HDAC1, MBD3, p65 and RBM14)**

[a] N-terminal GFP-tagged candidate proteins localize to nucleus. HEK293 cells were transiently transfected with N-terminal GFP tagged construct of candidate protein as indicated. Merged images show an overlap of DAPI and GFP signal (scale bar 50um). [b] N-terminal GFP-tagged constructs express the recombinant protein of expected molecular weight. Western blot analysis of extracts from HEK293 cells transiently transfected with N-terminal GFP tagged construct with GFP antibody.

**Supplementary Figure 4**

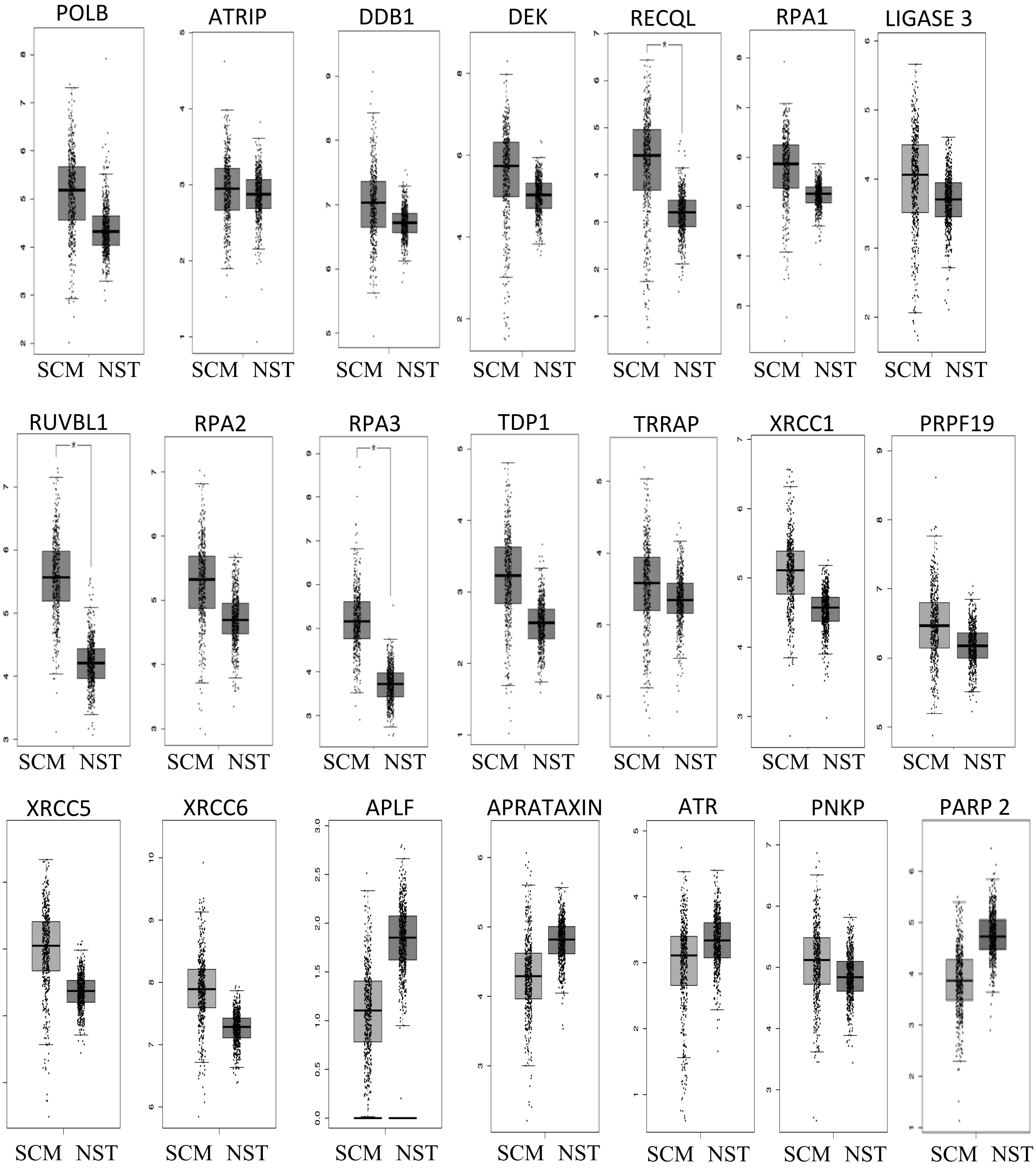


**Supplementary Fig. 4 GEPIA Analysis for expression data of DNA damage associated proteins in SCM vs. NST**

GEPIA is used to analyze the expression data of DNA-repair proteins that interact with vGAF, from 461 SCM and 558 NST samples. The expression levels of above mentioned genes were not found to be statistically different in two groups.
